# Assessment of Antibody Library Diversity through Next Generation Sequencing and Technical Error Compensation

**DOI:** 10.1101/085498

**Authors:** Marco Fantini, Luca Pandolfini, Simonetta Lisi, Michele Chirichella, Ivan Arisi, Marco Terrigno, Martina Goracci, Federico Cremisi, Antonino Cattaneo

## Abstract

Antibody libraries are important resources to derive antibodies to be used for a wide range of applications, from structural and functional studies to intracellular protein interference studies to developing new diagnostics and therapeutics. Whatever the goal, the key parameter for an antibody library is its diversity, i.e. the number of distinct elements in the collection, which directly reflects the probability of finding in the library an antibody against a given antigen, of sufficiently high affinity. Quantitative evaluation of antibody library diversity and quality has been for a long time inadequately addressed, due to the high similarity and length of the sequences of the library. Diversity was usually inferred by the transformation efficiency and tested either by fingerprinting and/or sequencing of a few hundred random library elements. Inferring diversity from such a small sample is, however, very rudimental and gives limited information about the real complexity, because complexity does not scale linearly with sample size. Next-generation sequencing (NGS) has opened new ways to tackle the antibody library diversity quality assessment. However, much remains to be done to fully exploit the potential of NGS for the quantitative analysis of antibody repertoires and to overcome current limitations. To obtain a more reliable antibody library complexity estimate here we show a new, PCR-free, NGS approach to sequence antibody libraries on Illumina platform, coupled to a new bioinformatic analysis and software (Diversity Estimator of Antibody Library, DEAL) that allows to reliably estimate the diversity, taking in consideration the sequencing error.

## Introduction

Antibody repertoires have been used in conjunction with display or selection technologies [1–7] and many libraries and antibody formats were created to satisfy the high demand for the different applications of recombinant antibodies [6,8–16].

The key parameter for an antibody library is its complexity [17], an estimate of the number of distinct elements in that collection. The amount of different functional species is directly related to the probability of that library to contain a functional antibody against a given antigen [18]. Despite the simplicity and the importance of this concept, until recently, measuring the diversity of antibody repertoires in a reliable and quantitative way was not possible and was approximated to the transformation efficiency of bacteria used to amplify the library [17,19,20]. To corroborate this estimate, so far the standard procedure in the literature consisted in testing the fingerprint pattern or the sequencing data of a few hundred library members for the presence of duplicates [14,21]. However, finding no identical clones in a random sample of a few hundred clones gives only a superficial evaluation of the library complexity and cannot be used to derive an estimate of the library complexity, which is expected to be from 10^4^ to 10^6^ times higher. The final complexity is then calculated by multiplying the estimated frequency of unique elements in the sample by the transformation efficiency. This calculation, albeit intuitive, is not correct, since the complexity does not scale linearly with the number of elements in the fingerprinted sample. The probability of finding a duplicated element grows as a function of the number of elements analyzed [22]. Indeed, the 10000th element does not have the same probability to be unique as the 100th element. Next generation sequencing has transformed functional genomics, and its application to the characterization of natural and synthetic antibody repertoires is growing [17,19,23]. The diversity parameter is however still hard to quantify precisely. The first widely used method in antibody library sequencing has been the Roche 454 pyrosequencing [17,24,25], that provides read lengths in the 300-400 bp range, suitable for antibody variable domains, but associated to a higher error rate (~0.5% per base [26,27]) and a lower throughput (10^4^ - 10^5^) [23] than other platforms. Other sequencing platforms, such as PacBio, can sequence up to 8500 bp but have a very low throughput (~10^4^) and a very high error rate [19,28]. High error rate is not a critical issue when there is an appropriate coverage and a suitable genome reference that allows errors to be corrected. In the study of antibody repertoires a full coverage is not yet feasible [29] and a genome reference is lacking by definition, because antibodies undergo imperfect genomic V(D)J rearrangement [19]. Indeed, if a discrepancy is found when comparing the library sequences, it is impossible to discriminate whether it has a biological origin or it is due to errors occurring in the processing of the sample (technical error) [30]. This problem is well known in the literature and different groups use different methods to address this issue [19,30]. DeKosky [31] uses a 96% percentage similarity criterion in sequence clustering, while Glanville [24] requires the sequence to have at least 2 amino acid mutations in at least one chain to be considered truly unique. Others [23,29] consider as reliable only sequences found at least twice or thrice in the sequencing step. We believe that such criteria are too strict to define the complexity, because a great amount of the sequencing data, that include the naturally occurring genomic modification, are eliminated. Moreover those methods neglect the error rate information derived from the sequencing data that can be used to resolve the discrepancy found.

In order to obtain a more reliable complexity estimate, we set up an Illumina-based sequencing strategy that, unlike previously described methods, is PCR-free, to avoid the PCR errors in the sample preparation. We then developed a software (DEAL (Diversity Estimator of Antibody Library)), which relies on base quality to solve the error rate problem. DEAL allows to get more accurate complexity boundaries increasing the sequence pool for the inferential estimate of the library complexity. We believe that the PCR-free sequencing and DEAL analysis could establish new standards for a more reliable and quantitative estimate of antibody library complexity and define quality criteria for newly created antibody libraries.

## Materials and Methods

### Construction of human SPLINT libraries

We constructed two SPLINT (Single Pot Library of Intracellular Antibodies) scFv antibody libraries from human RNA isolated from peripheral blood lymphocytes extracted from four anonymous voluntary donors which signed written informed consent, following a protocol modified from Marks and Bradbury [16] (see Supporting Information, S1 Fig). Blood samples were obtained from the Division of Transfusion Medicine and Transplant Biology, Pisana University Hospital, Pisa, Italy. IgM cDNA (μ heavy and κ and λ light chains) was used as a template to amplify VH and VL regions. Primers are designed to anneal to the external framework regions of the V genes. In the first library (hscFv1), each VH and VL subclass was first individually amplified, using every possible combination of the 5’ and 3’ primers available for VH and VL chains. The amplified products were then combined at equimolar ratios, so that each VH and VL subclass was equally represented in the library. In the hscFv2 library, instead, all VH and VL subclasses were amplified together in a single reaction (one for the heavy and one for the light V region) using a mix of the 5’ and 3’ primers available. A third single domain “nanobody” library (hVH) was created using only the VH domains, amplified in a single reaction. At the end of each library construction, the assembled VH-VL DNA products for hscFv1 and hscFv2, or the VH products for hVH, were ligated in the pLinker220 vector [32] for yeast expression in the SPLINT format, using restriction sites BssHII/NheI (NEB). Ligation of each library(~1μg) was transformed by electroporation into Max Efficiency E.coli DH5α cells (Invitrogen). Transformation efficiency for each library was assessed by colony count. See Supporting Information for details.

### Library DNA fingerprinting

100 individual bacterial colonies for each library transformation were picked and plasmid DNA extracted. Each scFv or VH present in the plasmid DNA was amplified by PCR and then digested with BSTNI (NEB) at 37°C for two hours. DNA fragments were resolved on a 8% acrylamide gel and stained with ethidium bromide. Pattern analysis was performed using Gel Quest software.

### Real Time PCR

Real time PCR was performed on the VH domain of the cDNA of each RNA sample used for library construction. All forward primers classes (BssHII-HuVHXaBACK, see Supporting Information, primers for VH) were tested against the most and the least abundant reverse primer class (HuJH4–5FOR and HuJH1–2FOR respectively). For each reaction, 4ng of cDNA sample were amplified following iTaq Universal SYBR Green Supermix (Bio-Rad) protocol. Real time PCR was performed on a Rotor-Gene Q Platform and the result analyzed following Nordgård et al. 2006 [
33].

### Sequencing sample preparation

To attach sequencing adapters to the scFv sequences, a ligation-based approach was designed. DNA adapters were synthesized harbouring overhangs complementary to the cleavage product of the restriction enzymes used for excising the scFv fragment from the plasmid, namely BssHII and NheI.

The forward and reverse strands of the adapters are synthesized independently and annealed in vitro (1:1 ratio, 95°C 5min, 95 → 25 °C in 5°C steps 1min/step). Before annealing the reverse strand was phosphorylated (0.2nmol Oligos, 10U PNK (NEB) 37°C 1h, 65°C 20min) to allow the ligation.

The scFvs were excised from the library plasmid (~8μg of the library were digested 3h 37°C with 4U of NheI (NEB), 3h 50°C with 4U of BssHII (NEB)) and ligated to the adapters (forward adapter: scFv: reverse adapter in 10:1:10 ratio, ~200-250 ng library 400U T4 ligase (NEB), O/N 16°C). The ligation was run on agarose gel and the band corresponding to the single insert with 5’ and 3’ adapters was resolved and purified with MinElute Gel Extraction Kit (Qiagen).

### NGS library processing

Libraries were quantified by Qubit dsDNA HS Assay Kit (ThermoFisher Scientific), diluted to 4nM, denatured with 0.1N NaOH (5' at RT), neutralized and diluted again in buffer HT-1 (Illumina) to a final concentration of 12.5 pM. Equimolar denatured Phi-X Control V3 DNA (Illumina) was spiked-in (20% of the volume) as an internal quality control and to increase the sample diversity according to Illumina guidelines. Sequencing was performed on MiSeq system with Reagent Kit v3 – 600 cycles (Illumina), with a cycle number of 350 and 250 for forward and reverse reads respectively.

Raw data were demultiplexed from .*bcl* files into separate .*fastq* files with *bcl2fastq-1.8.4* (Illumina), using the following barcodes as indexes: i1= TCAGCG, i2= GATCAC, i3= CTGAGA, i4= AGCTTT. In order to take into account the different length of shifter sequences introduced with the sequencing adapters, a specific number of nucleotides was discarded from the start of the reads (R1 index i1= 0, i2= 1, i3= 7, i4= 8; R2 index i1= 13, i2= 12, i3= 11, i4= 10). Reads were purged from adapter dimers, quality-filtered (Phred Score ≥ 32) and trimmed in sequences of the same length (R1: 320bp; R2: 220bp) *with trimmomatic-0.32* [34]. All the sequences whose forward and reverse reads both survived from the previous step were selected, taking advantage of the perl script *fastq-remove-orphans.pl*, which is part of *fastq-factory* suite (https://github.com/phe-bioinformatics/fastq-factory).

Forward and reverse-complemented reverse reads were then combined into 540bp-long pseudo-reads with a custom python script, and pseudo-reads from the 4 different indexes were pooled together. The hVH nanobody llibrary reads were merged using PEAR [
35], a pair-end read merger available at http://sco.h-its.org/exelixis/web/software/pear/.

### Software development

DEAL was developed as a standalone C++ program, without the need of external libraries. To manage the amount of data, a x64 compiler must be used during compilation. Speed optimization (MSVC: /O2 or /Ox, GCC: -O2 or -O3) should also be applied during compilation, to overcome tail-end recursion and to avoid long computing times. The program is tested to compile under Microsoft Visual Studio environment and work in Microsoft Windows 64-bit operating systems. DEAL was written in a modular structure to allow an easy future parallelization, even if not yet implemented. DEAL is available at 
http://laboratoriobiologia.sns.it/DEAL

Small complementary python scripts were created for format conversion, reads joining and primer recognitions.

R scripts were used for graphs generation and statistical analysis.

Pattern recognition for fingerprint analysis is implemented as a small Excel VBA script.

### Statistical analysis

The theoretical complexity of the hscFv1, hscFv2 and hVH nanobody libraries were estimated by the truncated Negative Binomial distribution (NBp,s, where p and s are the probability and size parameters) to fit number of sequences Nseq as a function of cluster cardinality x (see S2 Fig). Assuming Nseq(x) ~ C* NBp,s(x), it is possible to calculate the coefficient C, that represent the sought complexity to be estimated from the data (see details in Supporting Information).

## Results

### Library sequencing

Three sample libraries, 2 scFv libraries (hscFv1 and hscFV2) and a single domain library (hVH), were created from cDNA derived from human lymphocytes RNAs and amplified in bacteria (see Materials and Methods). Assuming that each transformed bacterium takes one copy of plasmid DNA, we can define the first hard cap of the library complexity as the number of total transformants obtained determined through CFU count. For the three libraries, hscFv1, hscFv2 and for the hVH nanobody library, this complexity upper bound is 15.8, 14.0 and 6.0 million elements respectively (Table 1). Standard fingerprint analysis on 100 random library clones for each library was performed and no duplicates were found (data not shown).

**Table 1.** Libraries complexity results.

| Library | Total reads<sup>a</sup> | Maximal complexity<sup>b</sup> | DEAL complexity<sup>c</sup> | Minimal complexity<sup>d</sup> | Theoretical Complexity<sup>e</sup> |
| --- | --- | --- | --- | --- | --- |
| hscFv1 | $6.02 \times 10^6$ | $15.8 \times 10^6$ | $3.45 \times 10^6$ | $0.6 \times 10^6$ | $8.96 \times 10^6$ |
| hscFv2 | $4.09 \times 10^6$ | $14.0 \times 10^6$ | $3.60 \times 10^6$ | $0.2 \times 10^6$ | $8.38 \times 10^6$ |
| hVH | $14.9 \times 10^6$ | $6.00 \times 10^6$ | $4.88 \times 10^6$ | $1.4 \times 10^6$ | $6.41 \times 10^6$ |

<sup>a</sup>Number of reads passing the first quality trimming analyzed by DEAL.
<sup>b</sup>Theoretical maximal complexity derived from the number of bacterial transformants.
<sup>c</sup>Number of DEAL clusters.
<sup>d</sup>Number of DEAL clusters minus the cluster with cardinality=1.
<sup>e</sup>Complexity estimated by truncated negative binomial fit.

### PCR-free sample preparation

In order to avoid any PCR-amplification step to attach sequencing adapters to the library to be sequenced which could insert mutations in the sequences, a ligation-based approach was designed (see Materials and methods). As shown in Fig 1, the forward read (R1) uses the P5 flowcell adapter (Illumina) and SBS3 sequencing primer (Illumina), while the reverse read (R2) uses the P7 adapter (Illumina) and SBS12 primer (Illumina).

**Fig 1.**
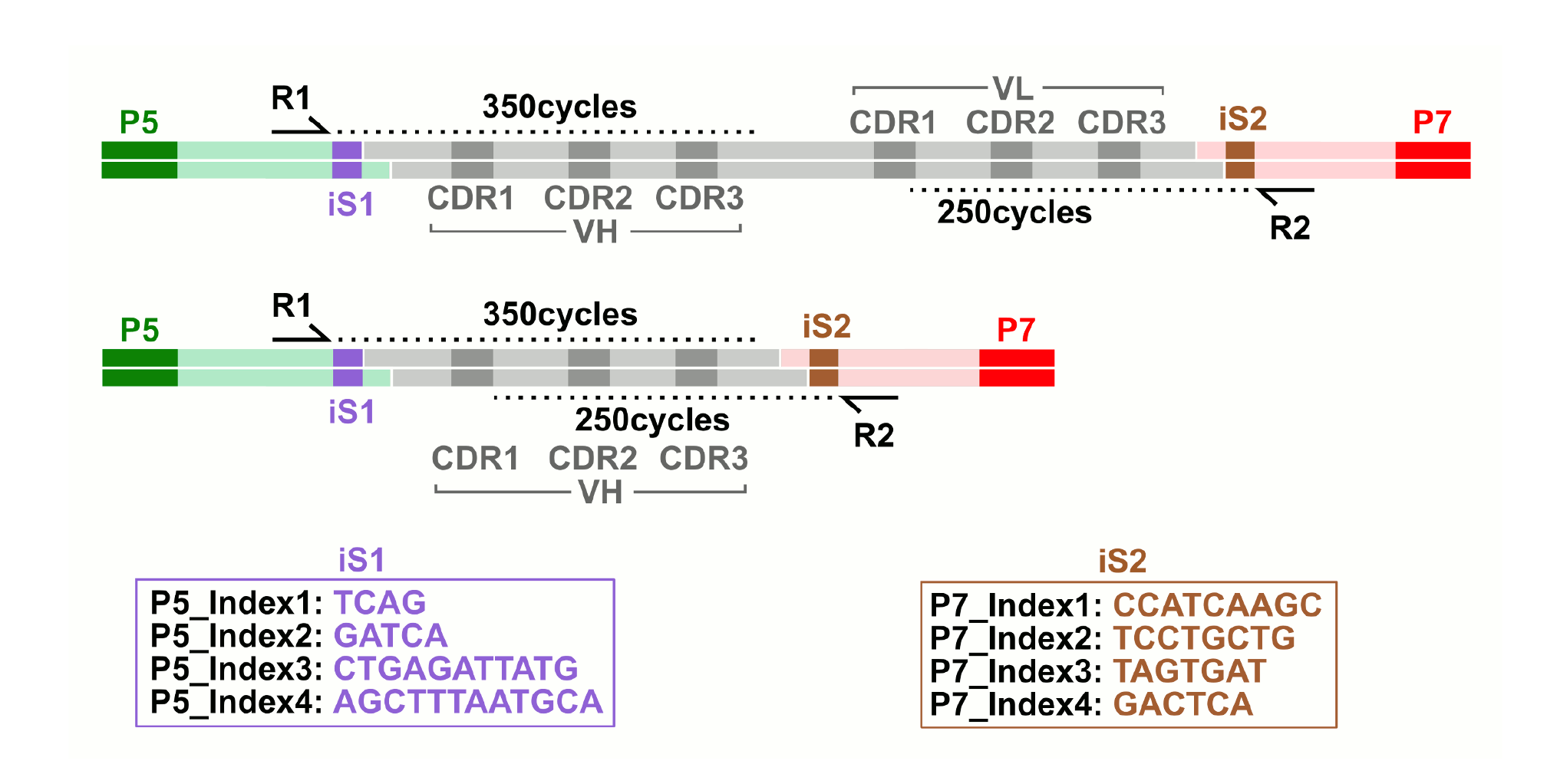
Diagram of sequenced adaptor-antibody-adaptor constructs. A) scFv library (gray), comprising heavy (VH) and light chain (VL) Complementary Determining Regions (CDR), was ligated to adapters (light green and pink) harbouring Illumina P5 and P7 flowcell hybridization sequences (green and red). B) VH nanobody library (gray), comprising heavy chain (VH) Complementary Determining Regions (CDR), was ligated to adapters (light green and pink) harbouring Illumina P5 and P7 flowcell hybridization sequences (green and red). The forward read (R1) uses SBS3 sequencing primer (Illumina), while the reverse read (R2) uses SBS12 primer (Illumina). iS1 and iS2 = index/shifter sequences.

Since sequences outside the Complementary Determining Regions (CDR), including the initial and distal framework regions, are rather constant, this poses serious technical limitations due to the sequencing by synthesis technology (SBS), which results in bad cluster recognition and low cluster count. To circumvent the problem, 4 barcoding index sequences (iS1 and iS2) were inserted just after each Illumina sequencing primer site. Using these barcodes allows to balance the read base content of the first 6 sequencing cycles. Moreover, inserting a different number of shifter bases before scFv fragment allowed to spread the eventual error in different positions of a sequence in the case of a error prone cycle, thus enabling correction during analysis.

### Sequencing

Libraries underwent SBS sequencing on Illumina MiSeq, executing 350bp forward (R1) and 250bp reverse (R2) reads after ligation of the adapters to the scFv fragments, excised by restriction enzyme digestion from the library vector (Fig 1; see Materials and Methods). 10-15 million of raw reads were produced for each experiment. The presence of the index/shifter sequence (iS) in the adapters successfully balanced the base composition of the first sequencing cycles, which are critical for cluster detection and run parameter estimation (S3A-C Fig). Median Phred score (Q-score) is an empirical measure of the confidence of base identification and remained greater than or equal to 30 (99.9% base accuracy) beyond the 300^th^ and 200^th^ cycle for R1 and R2 respectively (S3A-B Fig). Phred score is an estimate of the probability of error for any given base. The error rate distribution for hscFv1 library is shown in Fig 2A. On the other hand, the presence of a Phi-X phage DNA spike-in, as an internal control, allowed to assess the impact of sequencing errors on the library sequence analysis, by comparing Phi-X reads to the reference sequence, getting a real measure of per-tile and per-cycle error rates (Fig 2B and S3D Fig). Notably both the scale and the shape of these two distribution differ, leading to the conclusion that neither can be used alone for error rate estimation. While the median error rate was generally low (0.34±0.12%), both the presence of error rate “ spikes” (Fig 2B and S3D Fig) and the lack of homogeneous correlation between Phred score and the measured error rate (Fig 2C) revealed that particular attention must be spent, to distinguish the real base variants from background technical noise. The Phred-score-derived error rate and the Phi-X control error rate were similar in all the three libraries analyzed (data not shown). This data analysis revealed that the sequencing run was successfully performed. However, intrinsic technical limitations of the NGS, namely the presence of two unrelated error estimates requires an additional analysis. To this purpose a new software (DEAL) was developed.

**Fig 2.**
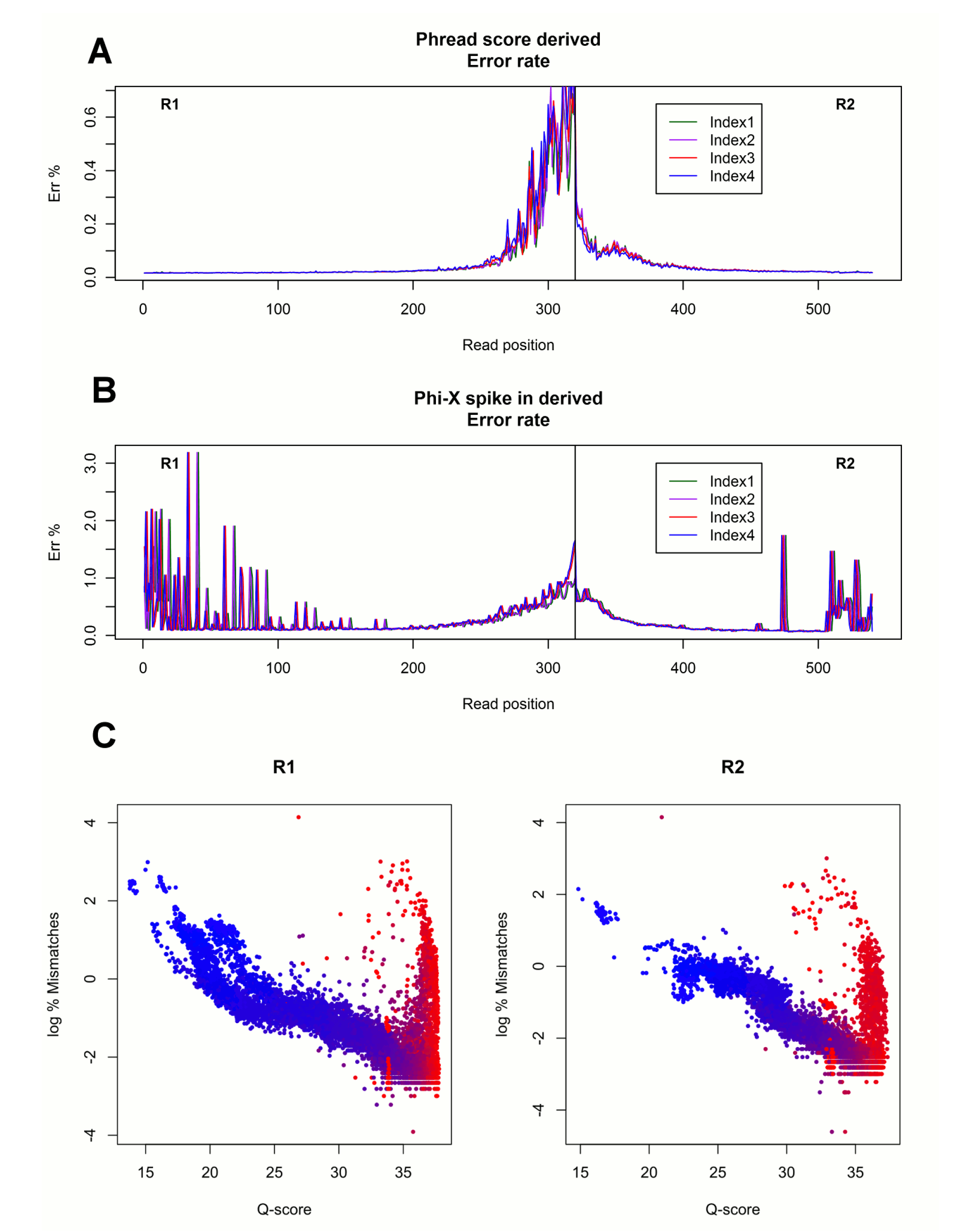
Phi-X derived and Phred score derived error rate distribution. A) Phred score error rate distribution for the hscFv1 library of the merged reads. Error rate increases with sequencing cycles. B) Control Phi-X derived error rate distribution for the hscFv1 library of the merged reads. Error rate is more prominent in the early sequencing cycles (spikes), with a small increase at the end of each read. The error distribution does not match the Phred score distribution and the shape differs as well. C) Scatter plot of the correlation of Q-score and log_2_(%Mismatches) in Phi-x control spike-in library. Each point represents the mean value from a single flow cell tile at a given sequencing read number, encoded by colour (red to blue: R1 cycle 1 to 350; R2 cycle 1 to 250; colour flex point is set at cycle 38). The Q score in the first 40 reads fails to be predictive of mismatch rate. Similar results were obtained for hscFv2 and hVH libraries.

### The Diversity Estimator of Antibody Library (DEAL) program

DEAL is a software tool to minimize the possible confusion between the real base sequence variants (biological diversity) from the background technical noise (technical misreading), taking into account both Phred derived error rate and Phi-X derived error rate.

DEAL is based on sequences identity collapse designed to skip the error prone bases. The number of the collapsed sequences is an estimation of the complexity of the analyzed library.

DEAL is divided in two main steps: in the first step, the sequences are clustered by identity, using a “ seed” of a 10-20 bp stretch in the CDR3s; in the second step, each element of the cluster is analyzed by binary comparison (Fig 3).

**Fig 3.**
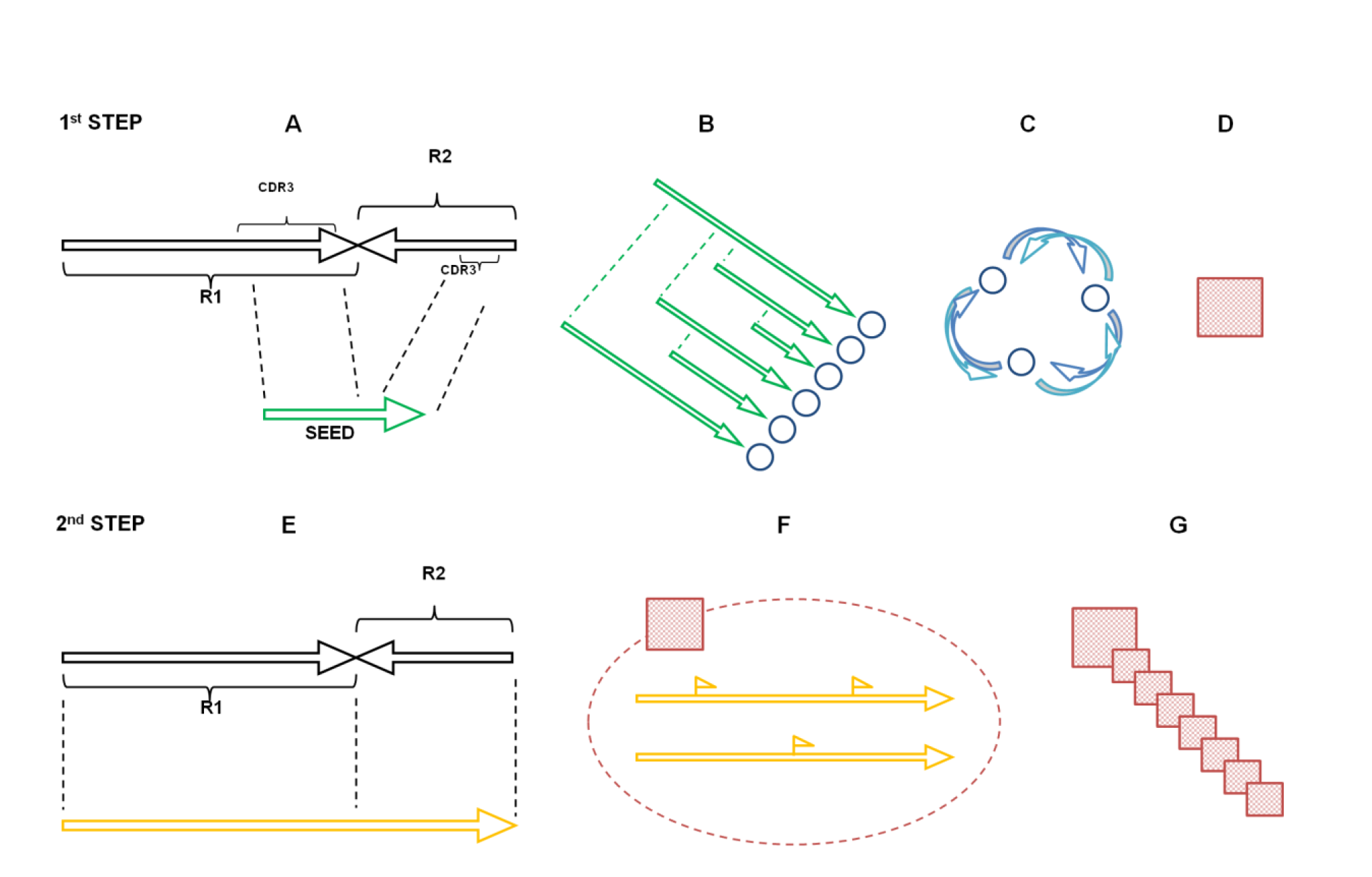
Diagram of DEAL workflow. 1^st^ STEP: A) Black arrows schematize the merged sequence, the green arrow represents the seeding region. B) The seeding regions are organized as a binary tree. Each seeding region has a sequence structure represented by the blue circle. C) During clustering, when two sequences are identical (they share the same branch of the tree), their sequence objects intertwine in a ring. D) At the end of the first step the rings of identical objects form a seeding group (red square). 2^nd^ STEP: E) Black arrows show the merged sequence, the yellow arrow represents the part of the sequence analyzed, by default the whole sequence. F) All the sequences in the same seeding group (i.e. have the same identical seeding region) are compared to each other for the whole length, mismatches (small flags) are taken into account. G) When two sequences in the group differ, a new subgroup is created (small squares). The total number of subgroups is the complexity of the analyzed library.

The first “seeding” step is necessary to reduce the calculation to local independent subproblems that can be solved by binary comparison. Moreover, this analysis is based on a system inspired by the crit-bit binary tree (Prof. DJ Bernstein http://cr.yp.to/critbit.html), which has the great advantage of being time and memory saving. The program works by creating a small sequence stretch from the seeding region provided and compares it to the tree of previous sequences. While confronting, in the presence of a difference, if it was not found before, a new branch is created (Fig 3B). If the position was already annotated, the sequence will be confronted to the corresponding branch. Each sequence has an associated sequence object. At the end of the comparison the identical sequences intertwine their sequence objects (Fig 3C). The first step ends when all sequences are analyzed and all the intertwined sequences form an individual cluster (Fig 3D). The number of these clusters is the seed diversity.

In the second step, the sequences within each cluster are compared along the whole sequence length (Fig 3E). The process is a binary comparison, so each sequence is compared to every previous analyzed sequence in its group. In this step, the possibility of sequencing errors is taken into account (Fig 3F). The program flags uncertain base read positions as unreliable, by checking both for a low Phred quality score in the sequences and for a high error rate in the sequencing cycles, that is retrieved from the error rate of control phage DNA (Phi-X). At this point, three scenarios can occur: i) if the sequences do not match, two subgroups are created; ii) if two sequences match and only in one an unreliable-flagged base is present, the base in that position will be assigned as the other sequence’s reliable base, and the two sequences are then merged and resolved as the same one; iii) if in two matched sequences at a specific position both bases are unreliable-flagged the resulting base in the merged sequence will be an unreliable-flagged base, which keeps track of the two possibilities.

The result of the computation is the creation of many small groups of matching sequences, whose number represents the sample complexity (Fig 3G).

Information about the grouping steps, as well as the final group distribution and all the sequences associated to the groups is also available using DEAL. It is possible to customize some parameters such as the seed positions and the optional computation of complexity of the deduced aminoacid sequence. In conclusion, DEAL software is able to ignore error prone bases during sequence identity collapse, leading to a reliable estimate of the library complexity.

### Upper and lower limit estimate of library complexity and outlier determination

The complexity of the three libraries was analyzed by NGS followed by DEAL. Data are summarized in Table 1. The sequencing cluster count (12.27, 10.98 and 15.52 million for human scFv libraries 1 and 2 and VH nanobody library respectively) was of the same order of magnitude as the upper cap complexity of the libraries, determined by transformants count (15.8, 14.0 and 6.0 million respectively). To be noticed, the VH nanobody library sequencing cluster count exceeded almost 3 times the upper cap limit, indicating an almost complete coverage of the library. Since it is crucial to have good quality sequencing data, the reads underwent a very strict quality trimming process. Only the reads which had a median Phred score of at least 32 (base call accuracy > 99.937%) survived the filter. The sequence count after trimming was 6.02 and 4.09 million for hscFv1 and hscFv2 respectively. A higher trimming survival count (14.9 million) was obtained for the hVH nanobody library, due to both the shorter length and the overlap of the two reads, which improves the median quality of the sequences.

DEAL was then applied to the quality trimmed data, using for the 2 human scFv libraries the default parameters: i) unreliable flag set when either error threshold for the Phi-X in the position was over 1% or quality in position is less than Phred 32; ii) seed position was located in the CDR3 of both VH and VL fragment (position 280-300 for read 1 and 470-490 for read 2). For the VH nanobody library, DEAL was set to allow a variable length in the input, the seed was placed in the 280-300 region only. Moreover the VH nanobody library protein complexity was also calculated.

The resulting distribution of clusters by cardinality is shown in 
Fig 4. A crowding in the first dozen groups can be seen, meaning that clusters with few elements are a clear majority. At the other end of the cardinality plot a clear single high cardinality element can be observed. This outlier element was present in all the analyzed libraries and was shown to originate from the backbone of the library plasmid vector, due to an incomplete digestion of the parental vector in the library construction process. In general the number of outliers is indicative of problems occurred during library construction, due for example to an overrepresentation of few specific clones. Indeed a high number of outliers would be indicative of an unbalance in the library lowering the chances of successful selections.

**Fig 4.**
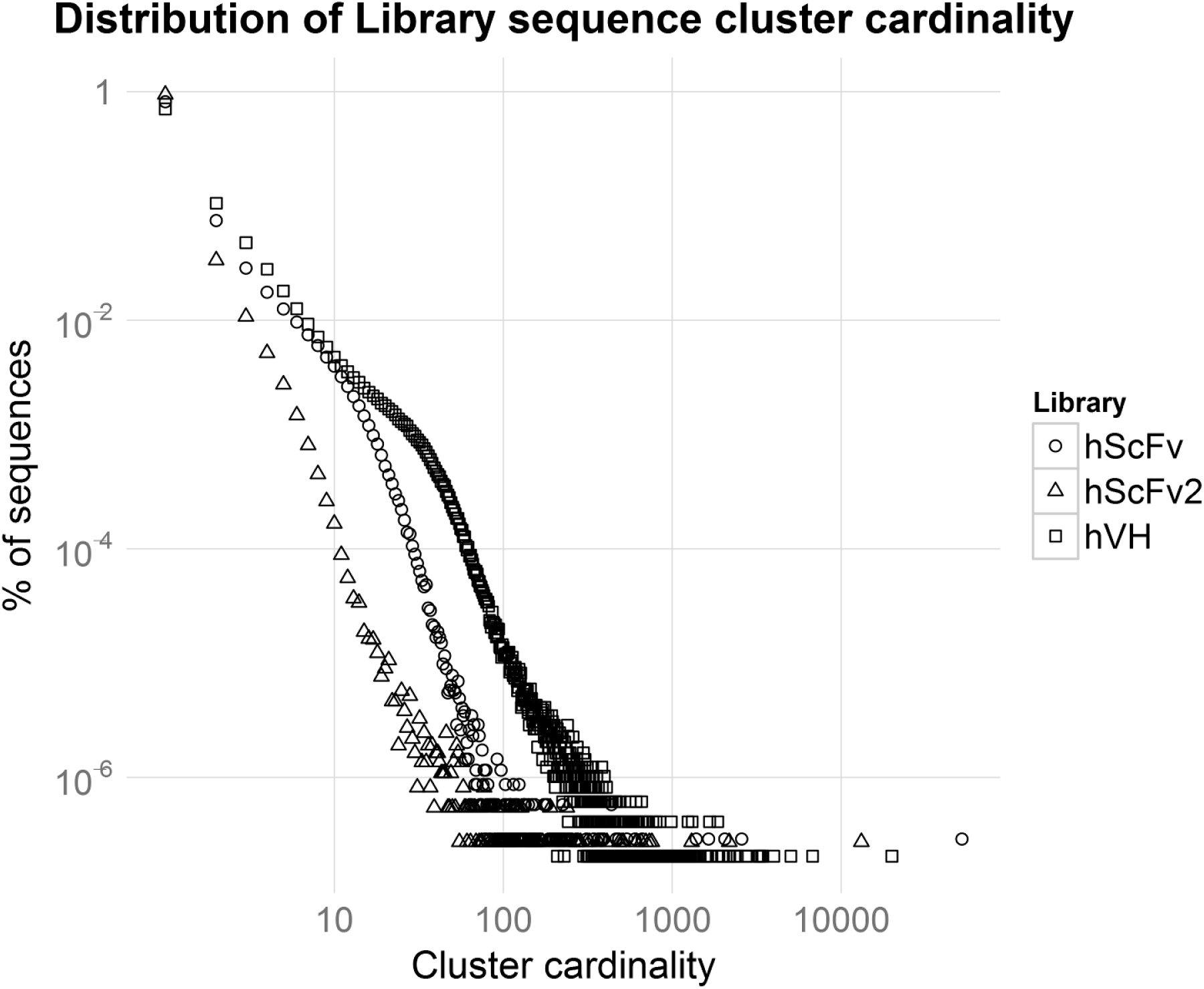
Distribution of library sequence cluster cardinality. Distribution of library sequence cluster cardinality. The more the curve is skewed towards high cardinality clusters, the lower the complexity of the library is expected to be.

Another kind of outlier is represented by the single element cluster (group with cardinality 1). This group contains, in addition to the real biological singletons (sequences with no other sequence in the same cluster), all the sequences where errors were not associated with a quality drop (and thus unable to be corrected by DEAL), that are unlikely to cluster with any other sequence in the sequencing run. For these reasons, the single element group is not reliable; therefore it should not be considered in complexity modeling to avoid an overestimation of the complexity. The number of DEAL clusters with cardinality two or greater represent the insurmountable lower limit of the library complexity (Table 1). The three libraries show respectively 0.6, 0.2 and 1.4 million unique clusters that satisfy the requisite, defining their lower limit complexity. However, if the coverage is not sufficient (i.e. for hscFv1 and hscFv2), raw data can not directly provide a measure of the total complexity. Thus a theoretical complexity calculation is also needed. To this purpose, an estimate of the theoretical complexity for each library was obtained using the truncated Negative Binomial distribution (Table 1 and S2 Fig).

### Library analysis: primer class distribution and VH-VL independent assortment validation

From the sequencing data, the chain independent assortment is another useful parameter to measure the skewness of a library. The NGS sequences were parsed with a custom python script for the primers used in the construction of the libraries. The distribution of the libraries by primer class is shown in Fig 5. A large portion of these are unclassifiable, probably due to the error spikes present in the primer regions, that impair the primer recognition, as discussed above (Fig 2B). The independent assortment of the VH-VL chains of the scFv libraries is shown in Fig 5A-B. No difference is observed between the frequency of forward and reverse primer pairs, compared to the theoretical combinatorial model, showing a good combinatorial assortment. Moreover, the frequencies of primers pairs assortment of the VH nanobody library matches the theoretical VDJ combinatorial model showing that very little amplification bias occurred during library construction (Fig 5C).

**Fig 5.**
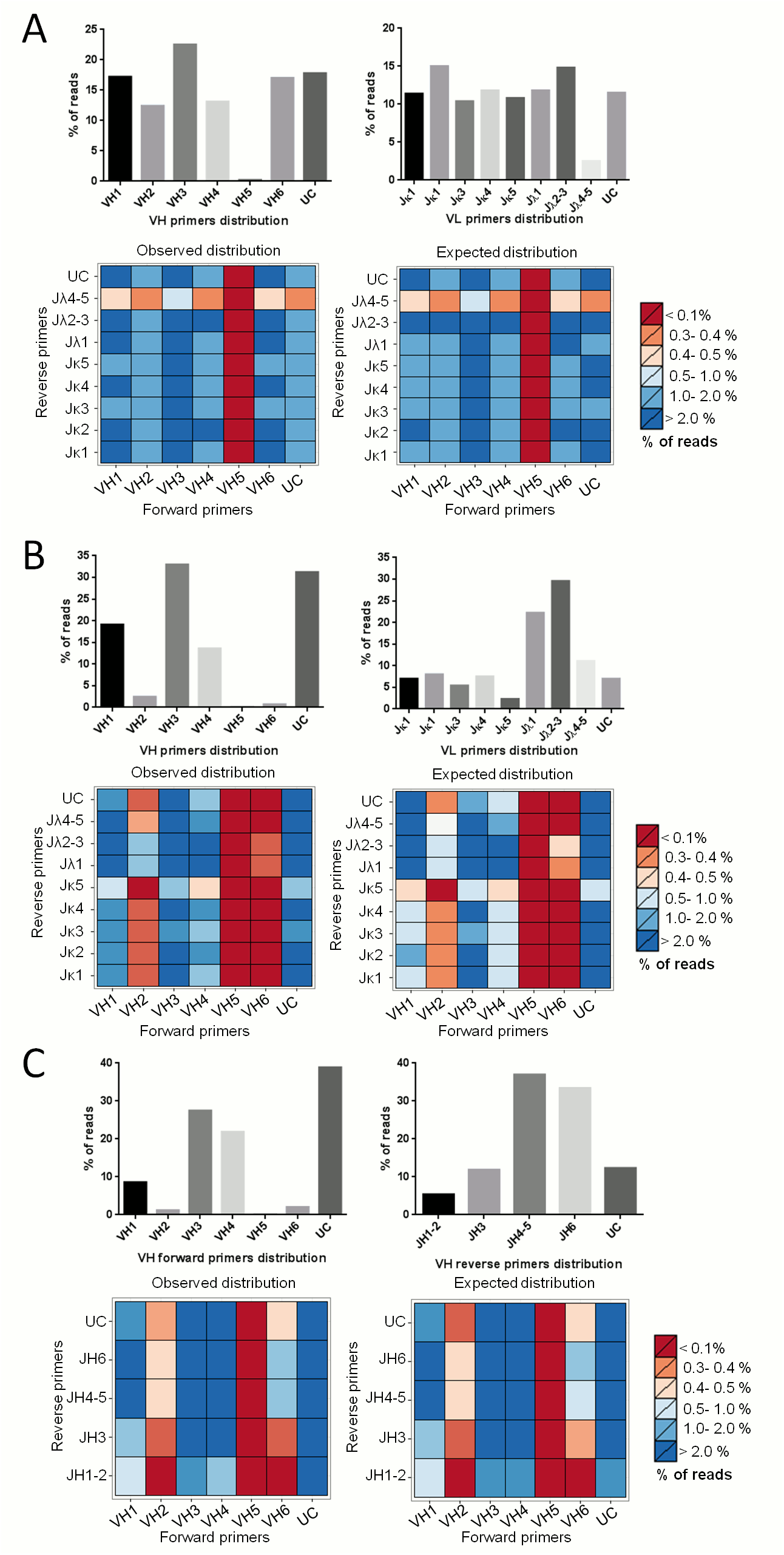
Chain/VDJ assortment independence of libraries. A) hscFv1. B) hscFv2. C) hVH. Top panels: barplots of forward and reverse primer distributions. Bottom panels: heatmaps of library primers distributions. Observed distribution is the primer pair proportion found after sequencing. Expected distribution is the multiplication of the two primers proportion (expected distribution given the independence between chains for the scFv libraries or given a balanced VDJ recombination for hVH). UC = unclassified. This category includes all the sequences that do not match any primer. The name of the primers is a shorter version of the original name listed in Supporting Information (Primer used for library construction).

The primers distribution, for all the three libraries, showed a very low percentage of reads for the class corresponding to the forward primer VH5 (HuVH5aBACK). To verify whether this reflected a problem related to NGS or to the real biological distribution, Real-Time PCR on the libraries and their corresponding cDNA was performed. Results showed that the same class distribution and abundance was present in the original cDNA, showing that the distribution bias of the VH5 primer class was not due to a sequencing issue (S4 Fig).

Usually, two different methods are used to amplify V regions for scFv libraries construction. In one, each VH and VL subclass is first amplified individually and then the products are combined at equimolar ratios [36,37]. Alternatively, all the VH and VL subclasses are amplified together in a single reaction (one for each type of V region) [16].

Therefore we compared the deduced complexity of two human scFv libraries, constructed by the two methods. To verify this issue, we constructed two human scFv libraries, using both protocols. The primer distribution obtained from NGS clearly reflects the two different approaches in library construction. hscFv1 shows a more poised distribution of all the V subclasses compared to hscFv2, in which, instead, there is a clear imbalance in classes representation (Fig 5). However, the Real Time data indicate that the protocol with the single common amplification reaction generates a library that faithfully reproduces the real distribution present in the natural repertoire. Thus, the first method guarantees an equal representation of all the different V subclasses, although it might not reflect the natural distribution.

### Determination of the protein diversity of a VH library through in silico translation of NGS reads

The quality of an antibody domain library is ultimately determined not only by its nucleic acid sequence complexity but also by the proportion of the sequences in the library that code for full length antibody domain protein. As the quality of a library resides in the protein conformational diversity, its most relevant estimator is protein complexity, which represents the true and relevant complexity of a library. While for scFv libraries the NGS reads do not yet allow to unambiguously deduce the full protein coding sequence by in silico translation, due to sequence length limitations, the shorter VH length allows the full coverage of the entire VH sequences, since R1 and R2 reads overlap in the central DNA stretch. Considering that the central region is read twice, if a sequencing error occurs at a given position of the forward read, it can be corrected comparing the base at the same position of the reverse read and vice versa. Thus, assigning that position to the base with the best quality between the discording pair, this correction lowers the global error rate and guarantees the best quality read in the critical CDRs regions and allows the correct frame to be determined, for amino acid sequence deduction. This is still not possible for scFv libraries, due to the undefined number of nucleotides present in the unsequenced gap between the VH and VL.

The protein diversity of the VH nanobody library was determined by in silico translation of DEAL clusters, where all synonym sequences were aggregated and both the frameshift and nonsense mutation were filtered out. The length distribution of the sequenced VH nanobodies shows that the correct frame is maintained in 86.57% of the analyzed sequences (Table 2, Fig 6). Since, in Illumina sequencing, the proportion of indel mutations is negligible, the frameshifts are mostly due to non-functional VDJ recombination. The majority of the sequences are around 370 nucleotide in length and the in frame sequences are the most abundant. The minority of out of frame (+ 1 or + 2 base pair) sequences shows an identical bell shaped distribution. Sequences with stop codons are also very rare: 10.3% considering the univocal stop codons in error-free sequences, 16% considering possible stop codons in error-flagged sequences. Indeed, filtered data shows that 80% of the hVH library sequences code for functional nanobodies (Table 2). The total protein diversity derived by DEAL clusters count is 4.73 million, which is reduced to 3.2 million by filtering the nonsense and frameshift incomplete peptides. Therefore, ignoring the protein group of cardinality 1 (due to presence of possible technical errors), the lower bound of the library complexity is 1.43 million proteins, of which 1.2 million are functional.

**Table 2.** hVH library protein complexity.

| Total reads<sup>a</sup> | % in frame<br>sequences<sup>b</sup> | % sequences<br>without stop<br>codon in<br>CDS<sup>c</sup> | % sequences<br>with full<br>length<br>CDS<sup>d</sup> | Total DEAL<br>protein<br>complexity<sup>e</sup> | Total DEAL<br>functional<br>complexity<sup>f</sup> | Minimal<br>DEAL<br>protein<br>complexity<sup>g</sup> | Minimal<br>DEAL<br>functional<br>complexity<sup>h</sup> |
| --- | --- | --- | --- | --- | --- | --- | --- |
| 14.9 x 10<sup>6</sup> | 86.57 % | 89.7-84.0 % | 83.4-77.9 % | 4.73 x 10<sup>6</sup> | 3.25 – 3.19<br>x 10<sup>6</sup> | 1.43 x 10<sup>6</sup> | 1.25 – 1.20<br>x 10<sup>6</sup> |

<sup>a</sup>Number of reads passing the first quality trimming analyzed by DEAL.
<sup>b</sup>Percentage of functional in frame sequences.
<sup>c</sup>Percentage of sequences which do not contain stop codons or do not contain possible stop codon in a error flagged sequence.
<sup>d</sup>Percentage of sequences in frame and without stops or possible stops codon.
<sup>e</sup>Protein complexity from DEAL protein clustering.
<sup>f</sup>Number of DEAL protein clusters with full length CDS.
<sup>g</sup>DEAL protein complexity minus the protein cluster with cardinality=1.
<sup>h</sup>DEAL protein complexity with full length CDS minus the protein cluster with cardinality=1.

**Fig 6.**
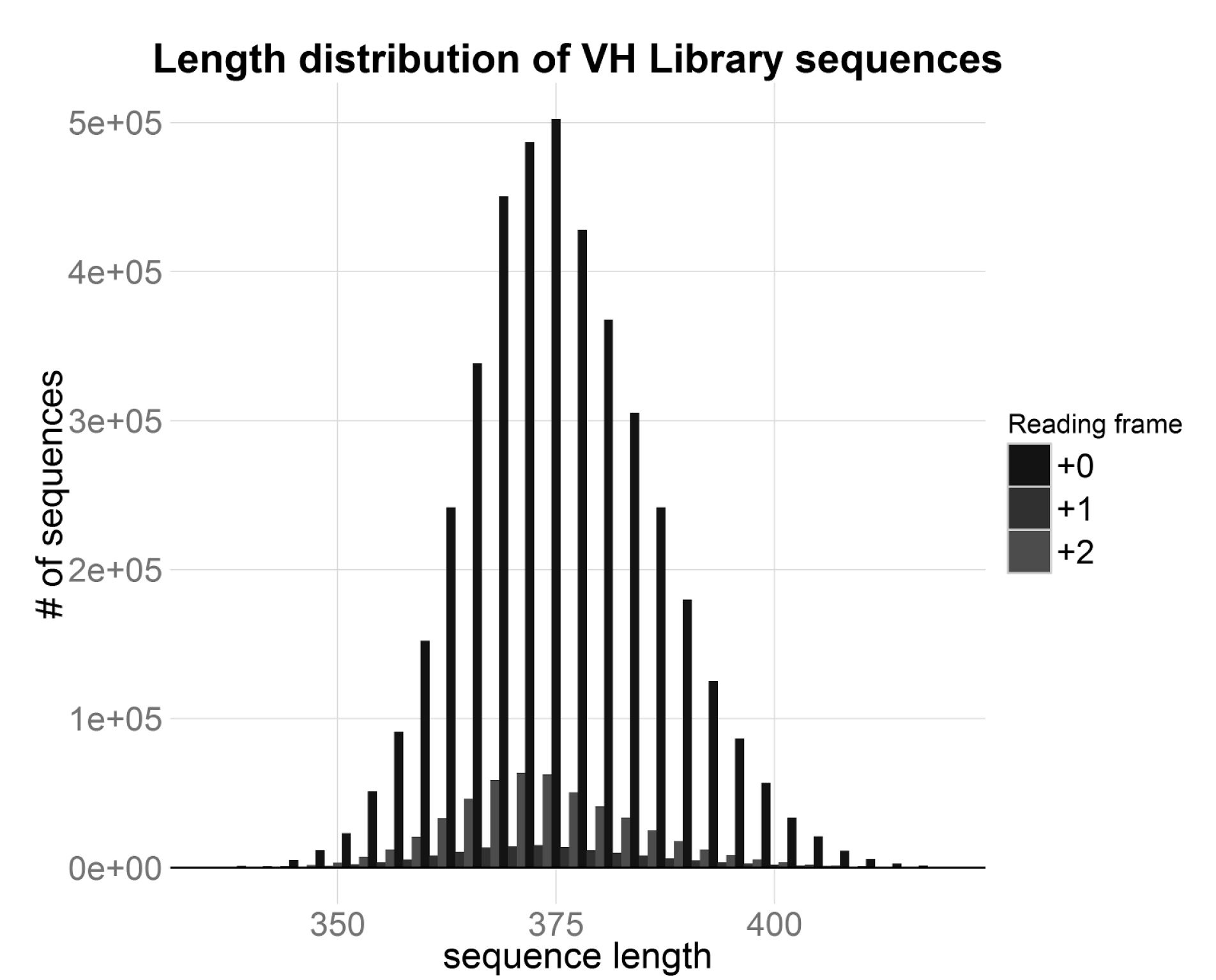
Length distribution of human VH nanobody library sequences. Barplot of the length distribution of human VH nanobody library sequences coloured by reading frame.

## Discussion

A quantitative evaluation of the sequence complexity of antibody libraries is critical to verify their quality and eligibility for screening purposes. Next generation sequencing is dramatically changing how antibody quality and complexity are estimated. We present here a PCR-free NGS sequencing strategy associated with the DEAL software, to compensate for technical errors, which allows to determine the minimal functional complexity of an antibody library. This method represents a significant advantage over current approaches. The main advantage of the proposed method is the definition of a “unique” biologically relevant sequence, filtering out as best as possible technical errors from the sequencing data. In fact, we found that Phred quality score generated by the sequencer alone is not sufficient to identify an error. Indeed, when comparing Phred quality to the error rate data extracted from the control phage DNA, it is clear that both parameters have to be taken into consideration.

In fact, Phred score is a pinpoint (both single sequence and single base) quality assessment, while the control phage DNA error rate can only be a chip-tile-wise quality assessment. Thus, it is not possible to substitute the Phred score with the error rate and both are required for the analysis.

In addition, to define the origin of an error we need to discriminate between pre-sequencing and sequencing errors. While sequencing errors need to be identified and removed (or limited), dealing with pre-sequencing errors is more complicated. In particular, some presequencing errors are introduced by the nucleic acid handling step during library construction (cDNA, V regions amplification). These errors are acceptable, sometimes deliberate [38], since they improve the diversity of the library. Instead, mutations introduced during the sequencing sample preparation (mainly PCR-derived) result in a misinterpretation of the sequences present in the library and thus have to be avoided. Therefore a PCR-free approach in the sequencing sample preparation is certainly to be preferred, since it reduces the number of errors (not associated to a quality drop) that DEAL cannot resolve.

The deduced amino acid sequence of the VH single domain library showed an unexpected length distribution, with more than 10.3% of non-functional out of frame sequences. Because in MiSeq indels are very rare, the length is most likely a natural feature rather than a sequencing artifact. As of today, the molecular mechanism whereby lymphocytes discard nonsense or out of frame variable domains and assemble only functional chains remains somewhat obscure. Interestingly, we observed in the hVH library a 7:1 ratio between functional and non-functional chains, which reflect the IgM RNA composition of PBLs. Indeed, this observation is in accordance with previous reports in mouse models [39,40]. In addition, the observed ratio appears to be remarkably high, even hypothesizing DNA silencing mechanism or RNA nonsense mediated decay taking place after transcription. In fact, if all the non-functional sequences were translated, there would be a considerable amount of useless transcripts and protein products. Moreover since this ratio refers only to a single chain, and an assembled antibody must have four correct chains to be functional, the ratio of functional assembled antibody would be even lower. Thus, it is clear that some kind of mechanism prevents the translation of these non-functional RNAs in vivo. Regarding the implications for antibody library screening, assuming the worst-case scenario where the light chains have the same functional proportion of the heavy chains, we could tentatively conclude that more than 80% of single chain library elements and more than 64% of scFv library elements encode a correct protein product.

Concluding, besides deep sequencing, the only other available method to determine the quality of an antibody library is a functional assay: the actual selection against an antigen. This is a valid practical strategy, although it is time consuming and can only give information on whether a library is adequate for screening purposes. The best general prediction of the capability of a library to undergo successful screenings is the estimate of its complexity in the most accurate possible way. Indeed, the complexity of a library can be directly linked to the probability to find a given binder of adequate affinity [18]. The method presented provides an advance towards this goal, by eliminating PCR steps and by compensating the technical errors. This method was validated for SPLINT intrabody libraries, but it is readily extendable to any antibody library of standard size. Moreover, DEAL software is independent of the sequencing platform and can be even more reliable with the greater number and more precise sequences that will be hopefully available in the future with the advancement of NGS technology.

